# Does the PolyJet Printing Technology Affect Mechanical Properties of 3D Printed Synthetic Tissue?

**DOI:** 10.1101/2024.06.18.599608

**Authors:** Samantha Kohnle, Emily Bermel, Leah Severseike, Varun Bhatia

## Abstract

Within the medical field, there are multiple opportunities in which 3D printed anatomical models could be utilized such as physician training, surgical planning, education, and R&D testing. Because of this, it is vital to choose materials that most closely mechanically mimic the tissues that would be seen within the simulated model. Stratasys currently has a line of Digital Anatomy (DA) printers that contain predetermined compositions and mixtures called anatomical presets to mimic soft tissue materials such as liver, myocardium, aortic, and epicardium tissue. Different compositions of these printed materials have been previously printed on the J750 Digital Anatomy printer and have undergone mechanical testing. Stratasys is releasing a new printer called J5 Digital Anatomy printer that will also have the capabilities of printing anatomical presets. To compare any differences between the two printer sample types, these mechanical tests were repeated on the J5 DA printer.

Mechanical testing included stiffness testing of printed liver samples, compliance testing of printed myocardium samples, and lubricity testing of printed cardiac tissue samples. Stiffness of J5 DA liver and J750 DA liver samples were similar with slight variation seen in more rigid J5 DA samples; however, both printer samples still fell within the stiffness range observed in porcine liver tissue. Compliance testing showed comparable stiffness between J5 DA and J750 DA myocardium samples; however, J5 DA samples tended to be slightly less stiff making them more similar to porcine myocardium stiffness. Thinner J5 DA samples were more variable when compared to J750 DA samples; however, this was minimal variability when compared to that experienced in porcine tissue. Lubricity testing showed comparable coefficients of friction between J5 DA and J750 DA samples with the exception of Mineral oil and Dry lubricants. J5 DA cardiac samples saw smaller coefficients of friction; however, these still fell within the range of aortic and epicardium coefficients of frictions. Because of these results, it can be concluded that the J5 DA printer samples are comparable to J750 DA printer samples and are alternatives to animal tissue benchtop testing depending on the application.

## Introduction

Recently, more interest in 3D printing has developed in multiple industries such as pharmaceuticals, aviation, defense, automobiles, architecture, entertainment, forensic, dentistry, audiology, medical, and food industry to develop products [1]. Within the medical field and industry, tissue mimicking models are highly sought after to aid in training, education, surgical planning, and quick prototyping purposes [2]. To create these phantoms, 3D printing is an efficient and cheap option to obtain specific anatomies or to simulate patient cases. Recent innovations within 3D printing such as improvements to bioprinting or the use of other material such as hydrogels have offered promising phantoms by creating whole organs that can be perfused and imitate similar bone and soft tissue properties found in medical imaging [3,4]. Depending on the intended application for the model, specific material properties may be needed to simulate tactile feedback. With continuously developing 3D printing capabilities, 3D printing has the potential to replace some cadaver and animal studies.

Previously, Stratasys (Eden Prairie, MN) has designed Digital Anatomy (DA) materials to simulate soft organ tissue and cardiac tissue within a J750 DA polyjet printer. Stratasys is introducing a new J5 DA printer that will have the anatomical preset printing capabilities. The J5 DA printer differs from J750 DA by printing on a small circular tray that rotates and moves up and down during the print process and is cured with a UV LED lamp while J750 DAP prints in a linear direction and is cured with a traditional UV lamp. Because of these differences, a comparative analysis was conducted between the DA samples from both printers and compared to porcine tissue. DA samples analyzed were DA soft organ tissue which is intended to simulate liver tissue and DA myocardium tissue which is intended to simulate the different cardiac tissue types. Mechanical testing on the new printer samples was repeated from the J750 DA material samples. These included DA liver types: Highly Contractile, Thicker Coated Highly Contractile, Moderately Stiff, and Thicker Coated Moderately Stiff. Myocardium DA printed samples included were Highly Contractile, Very Stiff and Extremely Stiff. The repeated mechanical testing included: i) mechanical compression testing of DA soft organ samples to compare to porcine liver ii) mechanical compliance testing of DA Myo to compare to porcine myocardium found in the right atrium, right ventricle wall, ventricular septum, left atrium, and left ventricular wall, and iii) lubricity of DA Myo compared to porcine epicardium and aorta.

## Methodology

### Porcine Tissue Testing

Porcine tissue testing was previously conducted in prior mechanical testing and was used to compare to 3D printed samples [5,6]. Porcine tissue was used to compare the printed tissues due to this being a common pre-clinical animal model. All three mechanical tests: compression, stiffness, and lubricity were performed on porcine samples obtained from the liver, right atrium, right ventricle, ventricular septum, left atrium, and left ventricle. This tissue was collected from butchers and pre-clinical research facilities and frozen at -80C either immediately after termination or 2-4 hours after termination. Samples were allowed to thaw in an insulated cooler prior to testing for around 14 hours [5,6].

### Stiffness Testing

A uniaxial Instron with a 50N load cell was used to perform the stiffness test (compression). A 9Fr round rod simulating a surgical tool was mounted to three-jaw chuck applied an axial compression at a rate of 0.5in/min. Liver DA printed samples were placed in a rigid 3D printed test fixture with a bottom to allow for the sample to compress similarly how it would in vivo (Fig 1). Cubic 3D printed samples with dimensions of 30×30mm with varying thicknesses of 15mm and 25mm were tested. Each liver type and thickness combination were contained a sample size of six.

**Figure 1.**
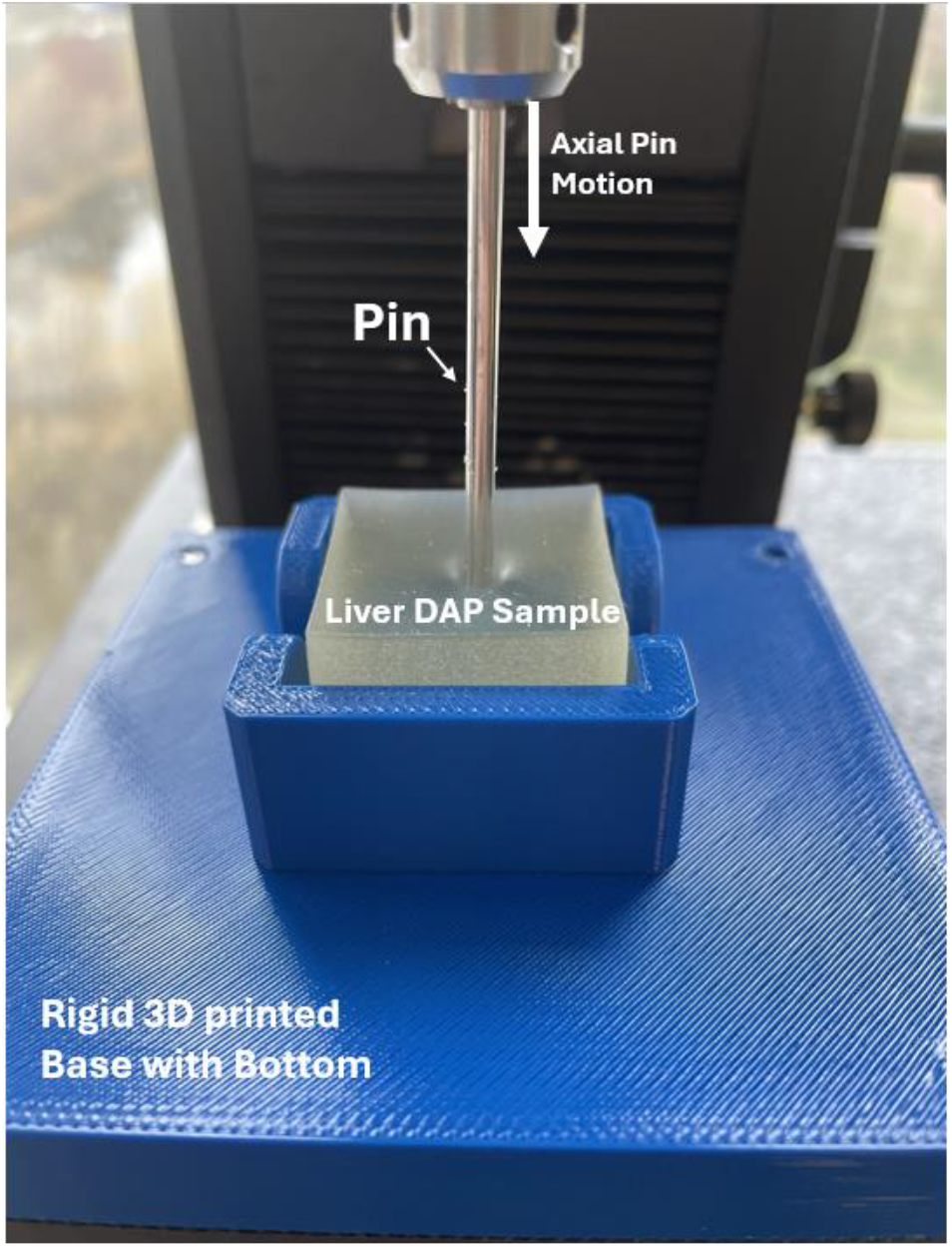
Printed Liver sample showing compression from a 9Fr rod during the stiffness test

After data collection, a force vs distance graph was generated where stiffness was determined by the slope of the graph. To account for safe and normal use conditions experienced by surgical tools and eliminate noise, stiffness was calculated between 1-3N.

### Compliance Testing

A uniaxial Instron with a 50 N load cell at an axial compression rate of 0.5in/min was used in the myocardium compliance tests. The printed myocardium sample was secured with screws in between two plates with a 1 inch diameter hole cut out (Fig 2). This was used to better mimic the boundary conditions that may be experienced in cardiac anatomy. Two different sized rods (9Fr and 22Fr) were used for the compliance test to mimic different cardiac delivery tool systems such as the smaller cardiac lead delivery systems and larger device delivery systems such as Medtronic’s leadless pacing system. Three myocardium printed sample types were used: Highly Contractile, Very Stiff, and Extremely Stiff. Cylindrical samples were printed with 2in diameter and two thicknesses, 2.5mm and 12.1mm. These thicknesses were chosen previously to correlate to different areas of cardiac anatomy that are thinner such as the atrium and ventricle free wall or thicker areas such as the ventricular septum. Each fixture rod size, myocardium type, and sample thickness combination contained a sample size of 12.

**Figure 2.**
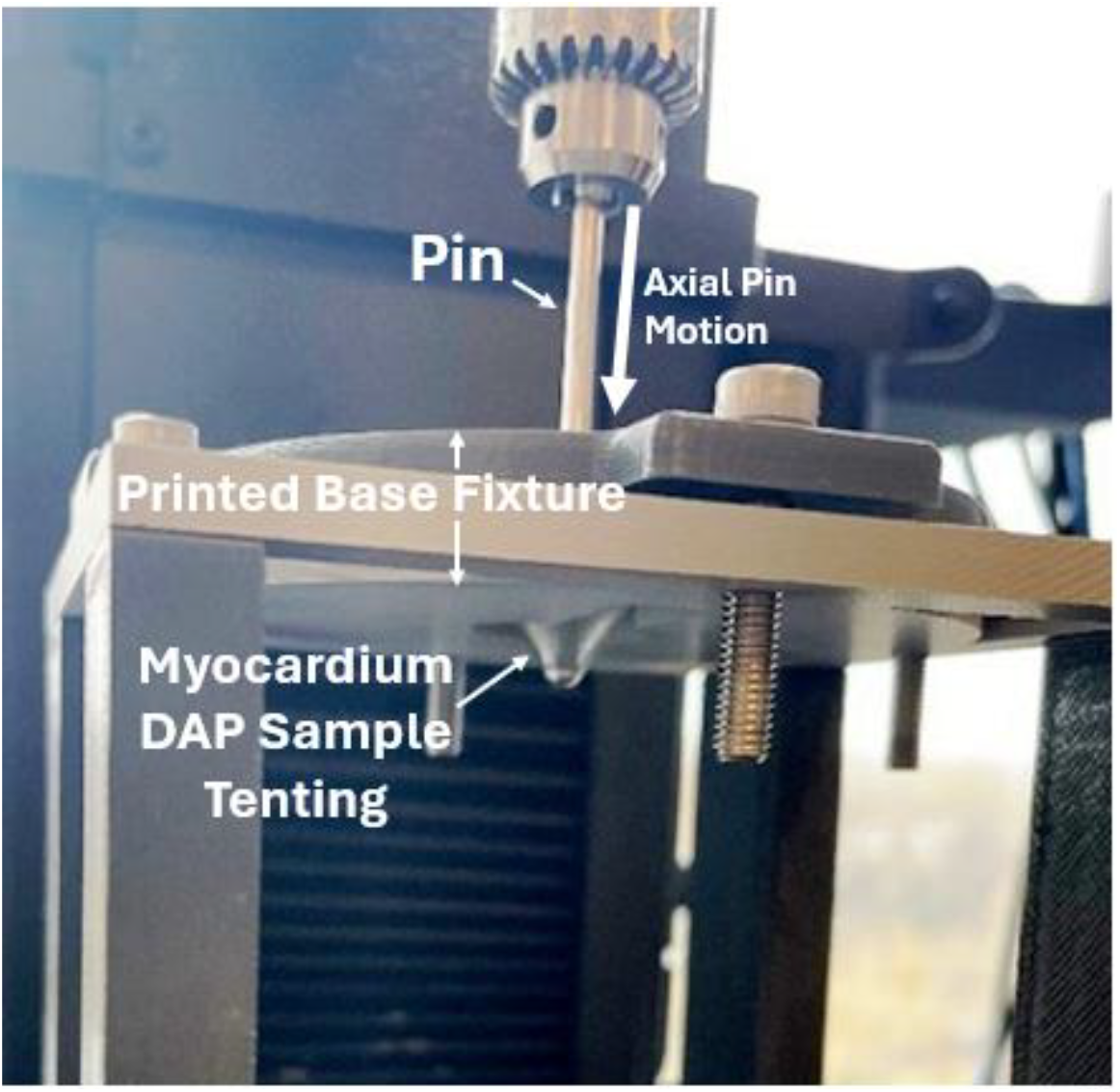
Printed myocardium sample (2.5mm) showing tenting from a 9Fr rod penetration

Data was processed and analyzed similar to how the previous iteration samples printed on the J750 printer were completed. A preload of 1N was applied during processing, the slope was examined as the sample was displaced until failure. The preload of 1N was chosen to eliminate potential noise that would be experienced by use conditions such as trabeculae or other cardiac anatomy. The slope of best fit was recorded in two displacement zones, 0-5mm and 5-10mm after the preload (Fig 3). Displacements of 0-10mm were used to analyze printed tissue stiffness due to use conditions experienced during cardiac implant procedures.

**Figure 3.**
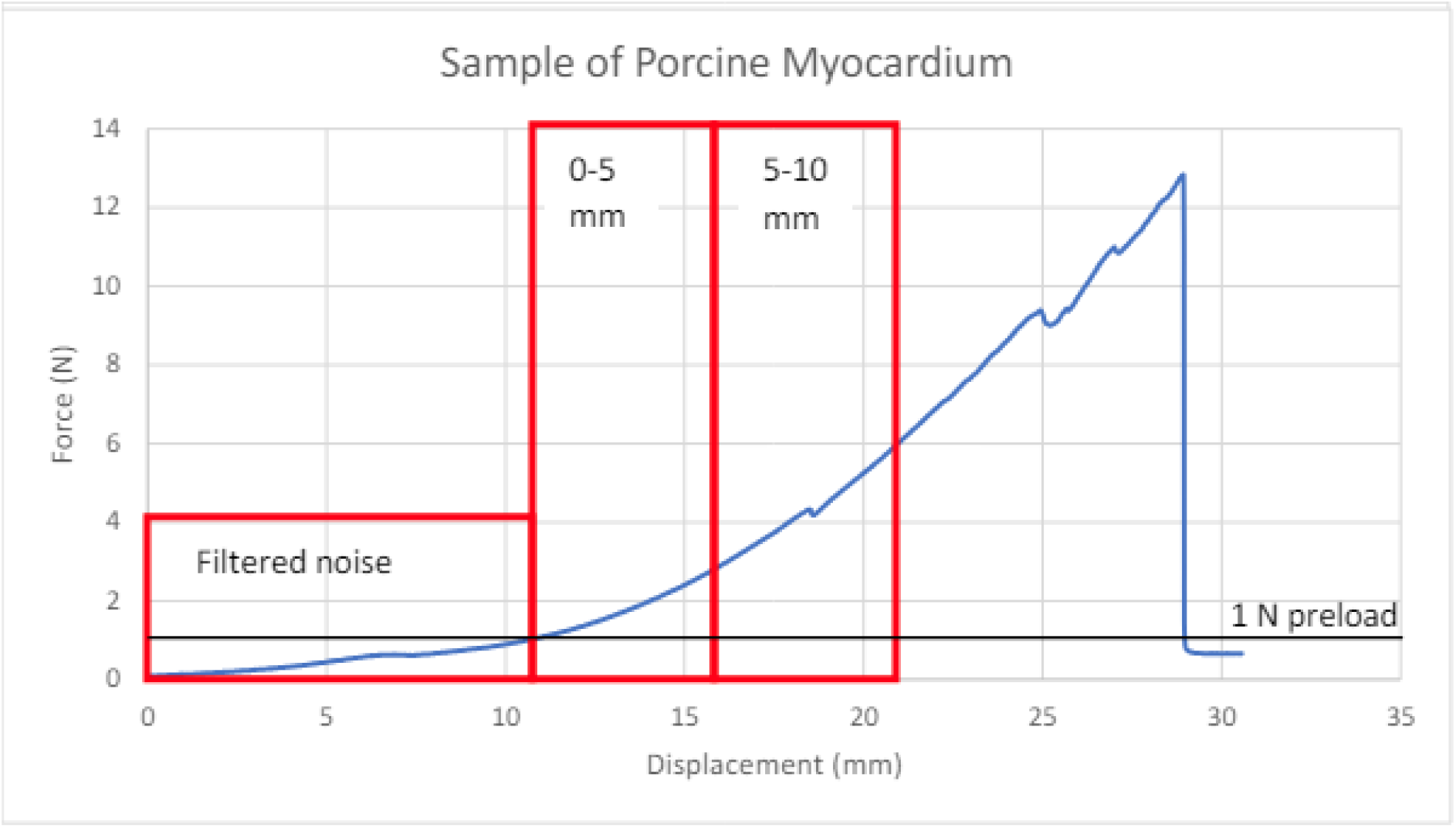
Example of raw data processing to determine the stiffness from 0-5mm displacement and 5-10mm displacement after filtering noise from a 1N preload [5]

### Lubricity Testing

A range of lubricants including no lubricant, DI water, soap, and mineral oil were analyzed on three DAP myocardium types, Highly Contractile, Very Stiff and Extremely Stiff. Each myocardium sample was printed with 10c10×0.6cm dimensions and a sample size of six was tested for each DAP myocardium type and lubricant combination. Lubricant samples received a thin layer (1-2ml) of designated lubricant type prior to testing. A 0.25in diameter steel ball probe was fixated to an Rtec (San Jose, CA) tribometer. An axial force of 0.75N was applied at a velocity of 0.5mm/s across 30mm while sampling every 0.1s. A rigid 3D printed test fixture held that sample in place as the fixturing head moved laterally as seen in Fig 4.

**Figure 4.**
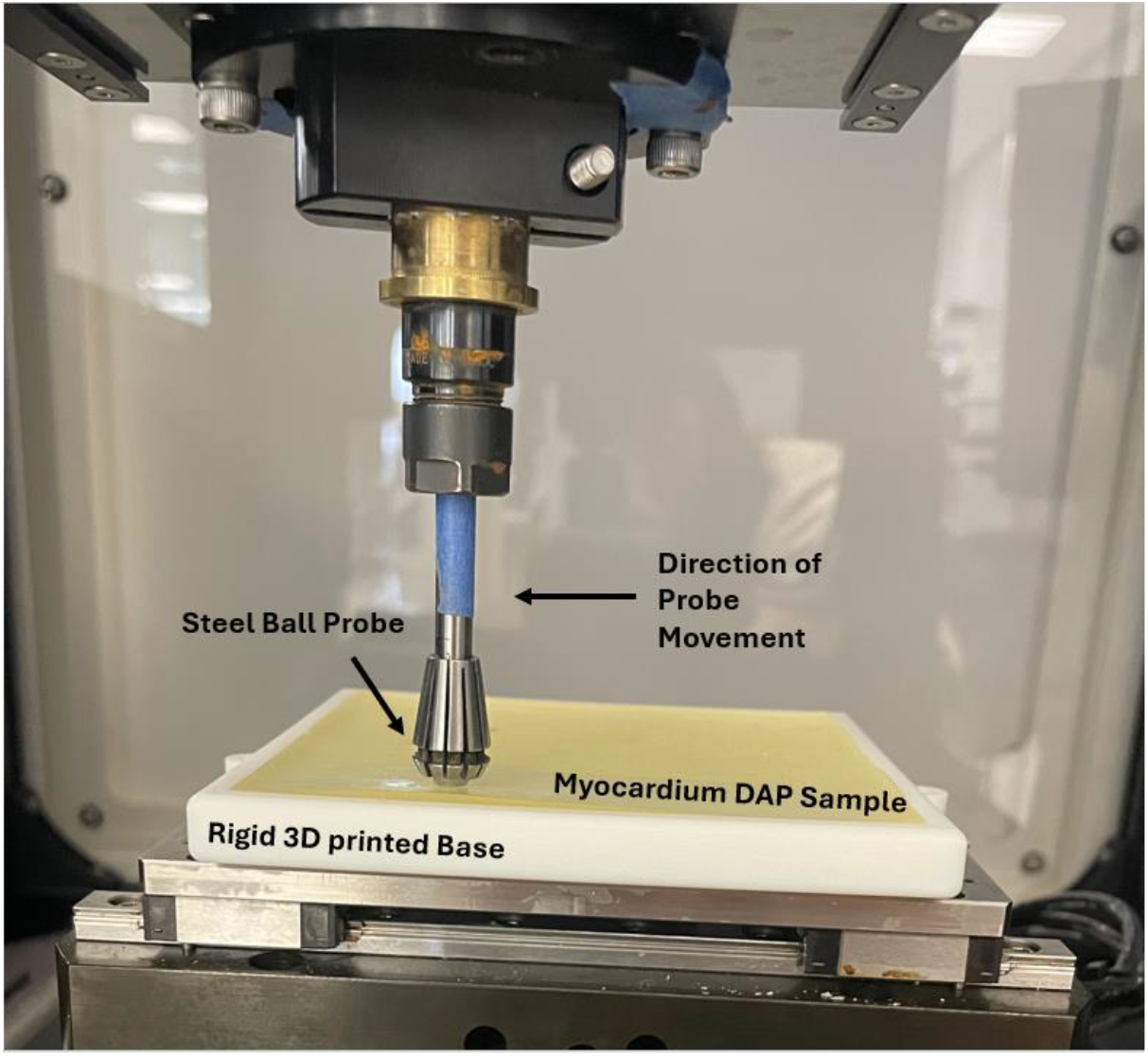
Lubricity test fixturing showing the printed myocardium sample loaded into the rigid 3D printed test fixture. The rod travels from the middle of the sample to the left for each iteration.

The raw data was noisy due the sensitivity of the load cell and low load conditions needed for compliant materials. To avoid any ramp up and down regions across the sample, the data was analyzed between 10-50s. Similar to the previous mechanical tests, a simple moving average was applied to the raw data and the coefficient of friction was recorded seen in Figure 5 [6].

**Figure 5.**
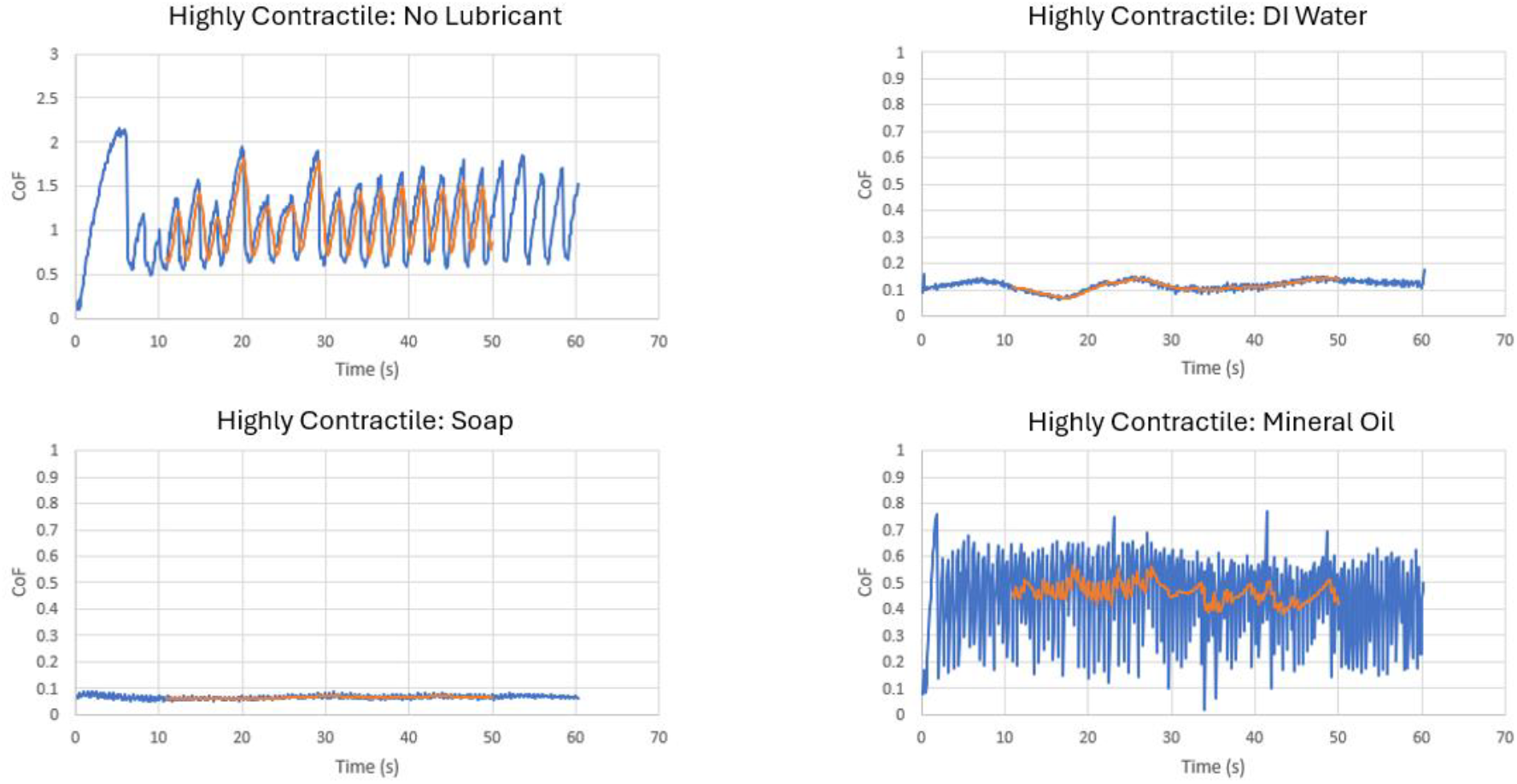
An example of the raw lubricity data shown in blue and simple moving average in orange between different lubricant applications

## Results

This characterization testing was performed with limited sample sizes to gain an understanding of general comparison between sample types. There were not enough samples tested to allow for full statistical comparison between J5 DA samples, J750 samples, and porcine tissue.

### Stiffness Testing

All printer liver type stiffness values fell within the range of porcine liver stiffness values (Fig 6). Moderately Stiff and Thicker Coated Moderately Stiff from the J5 DA printer saw greater stiffness values in both thicknesses compared to J750 printer samples and Highly Contractile from the J750 printer showed lower stiffness values between both thicknesses comparatively.

**Figure 6.**
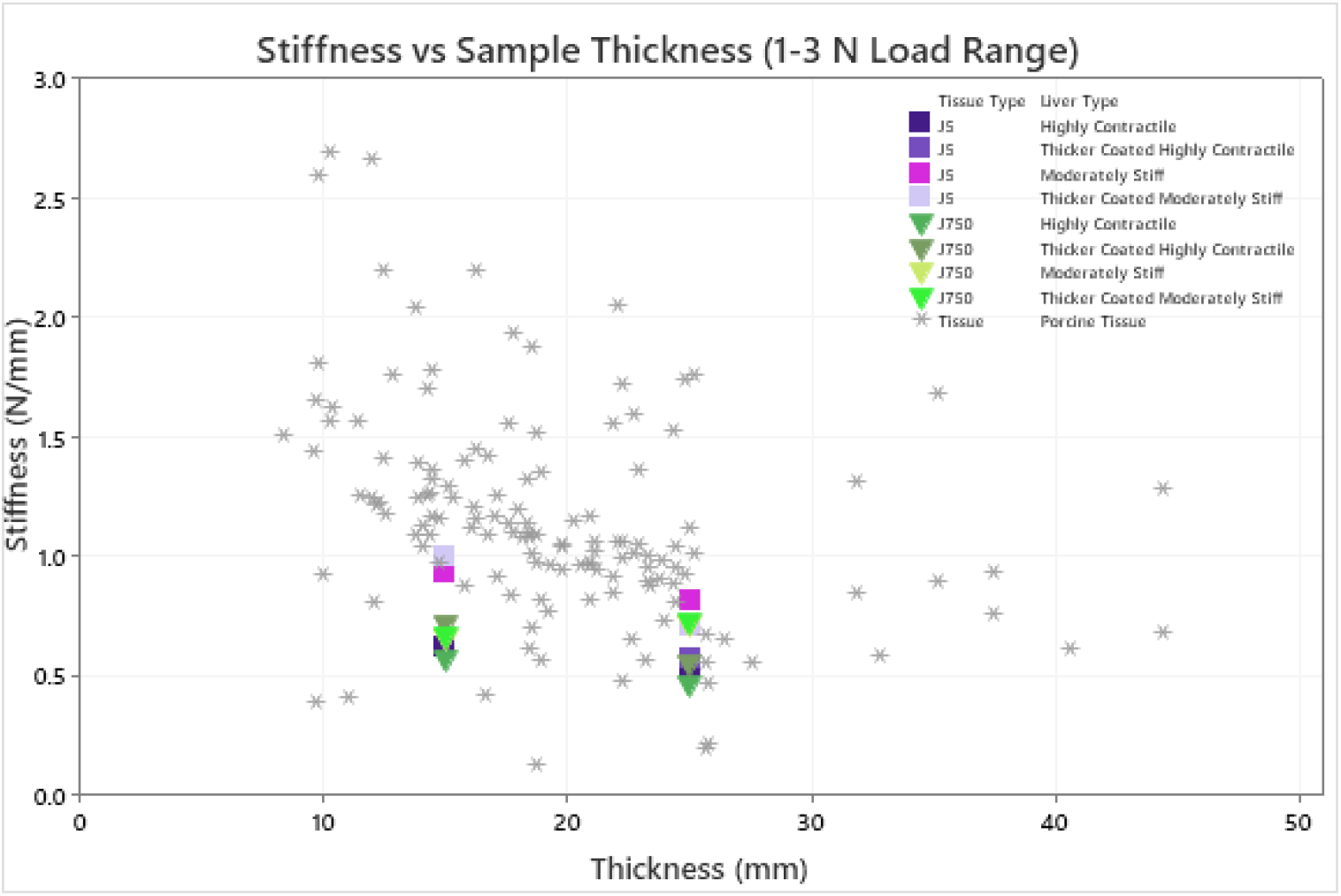
J5 DAP and J750 DAP liver samples at two thicknesses compared to porcine liver samples of variable thicknesses from multiple liver lobes

In less compliant samples such as Moderately Stiff and Thicker Coated Moderately Stiff at lower thickness values, J5 DA printer samples saw higher stiffnesses when compared to J750 DA printer samples. Higher compliant liver samples, Highly Contractile and Thicker Coated Highly Contractile, had comparable stiffness values between printers. For thicker samples, Highly Contractile, Thicker Coated Highly Contractile, and Thicker Coated Moderately Contractile had comparable stiffness values between printers whereas Moderately Stiff saw slightly higher stiffness values in J5 DA printer samples (Fig 7).

**Figure 7.**
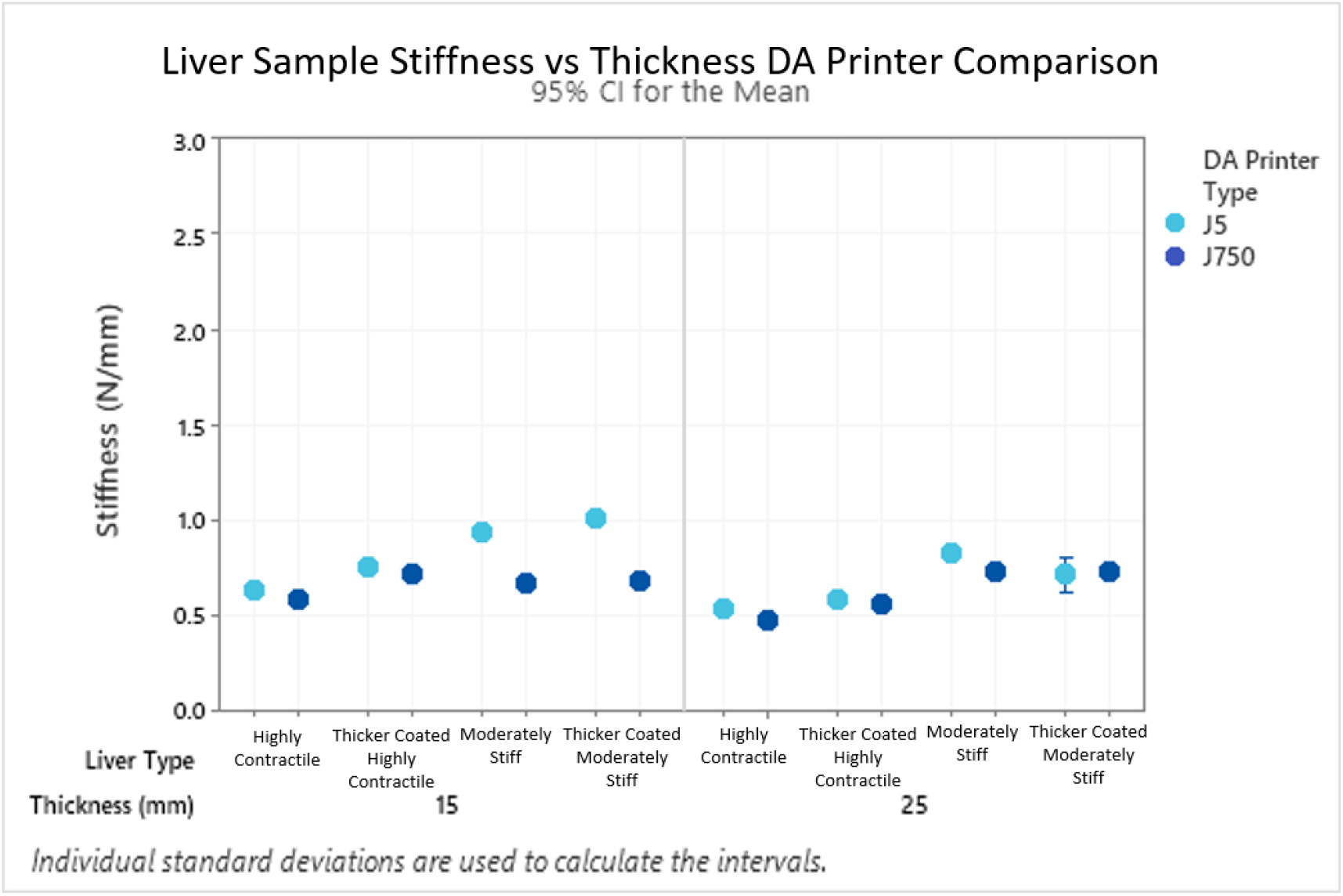
Stiffness of liver compression tests comparing two printer type samples

### Compliance Testing

Samples that underwent small pin stiffness testing at a displacement of 5mm didn’t observe major differences between printer types at different sample thicknesses. J5 DA printer samples were within the stiffness range observed from various porcine cardiac locations (Fig 8).

**Figure 8.**
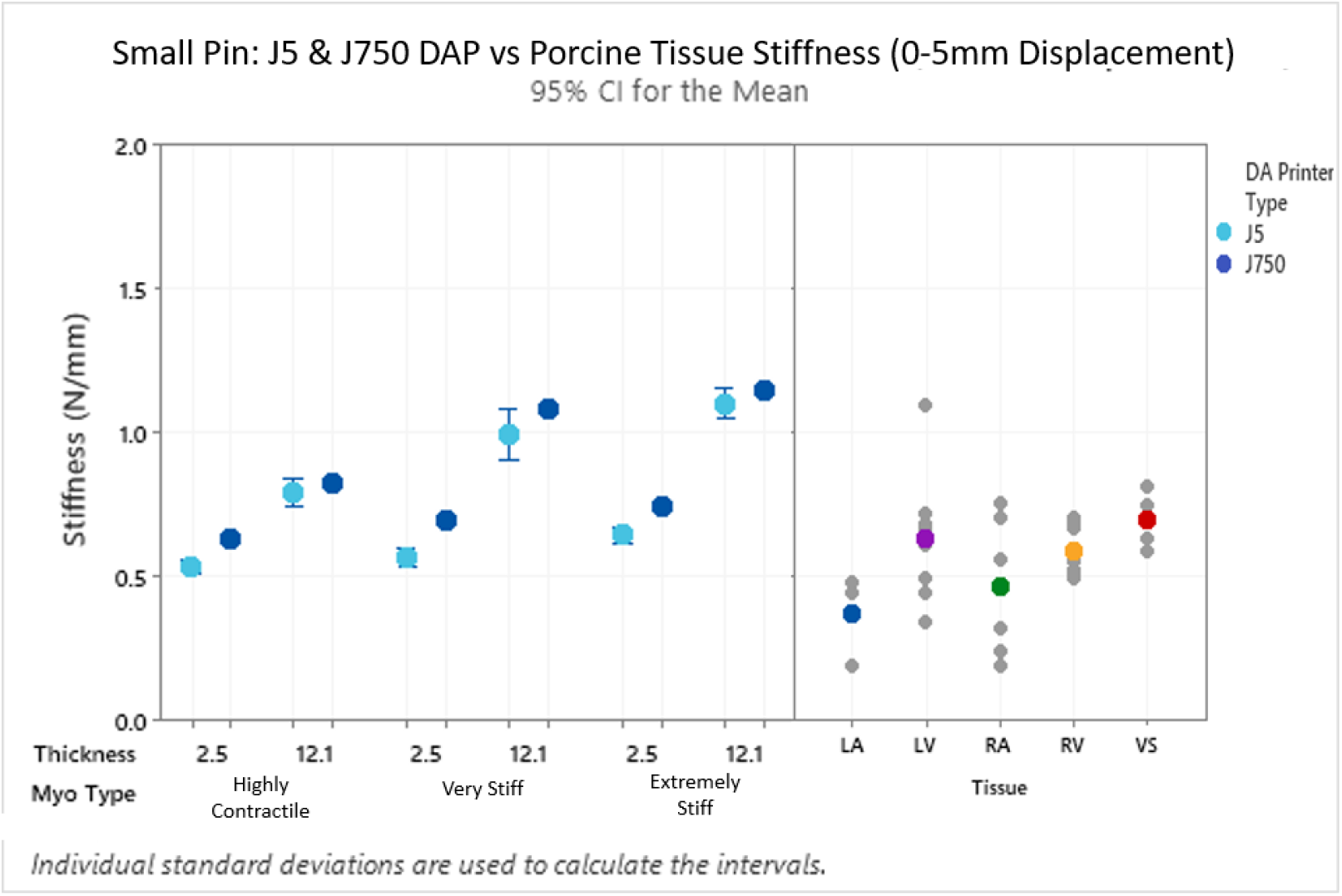
Stiffness comparison of the J5 DA Printer and J750 DA Printer myocardium samples compared to porcine tissue using a 9Fr pin displaced within the first 5mm.

With a small pin displaced 10mm, thin samples from the J5 DA printer saw more variability between stiffnesses when compared to the J750 DA printer; however, these averages still fell within the porcine cardiac tissue ranges. Between the J5 DA and J750 DA printed samples, J5 DA samples were less stiff between thickness and tissue types (Fig 9). Because of this, these stiffness values are then more comparable to stiffness values observed in porcine cardiac tissue samples.

**Figure 9.**
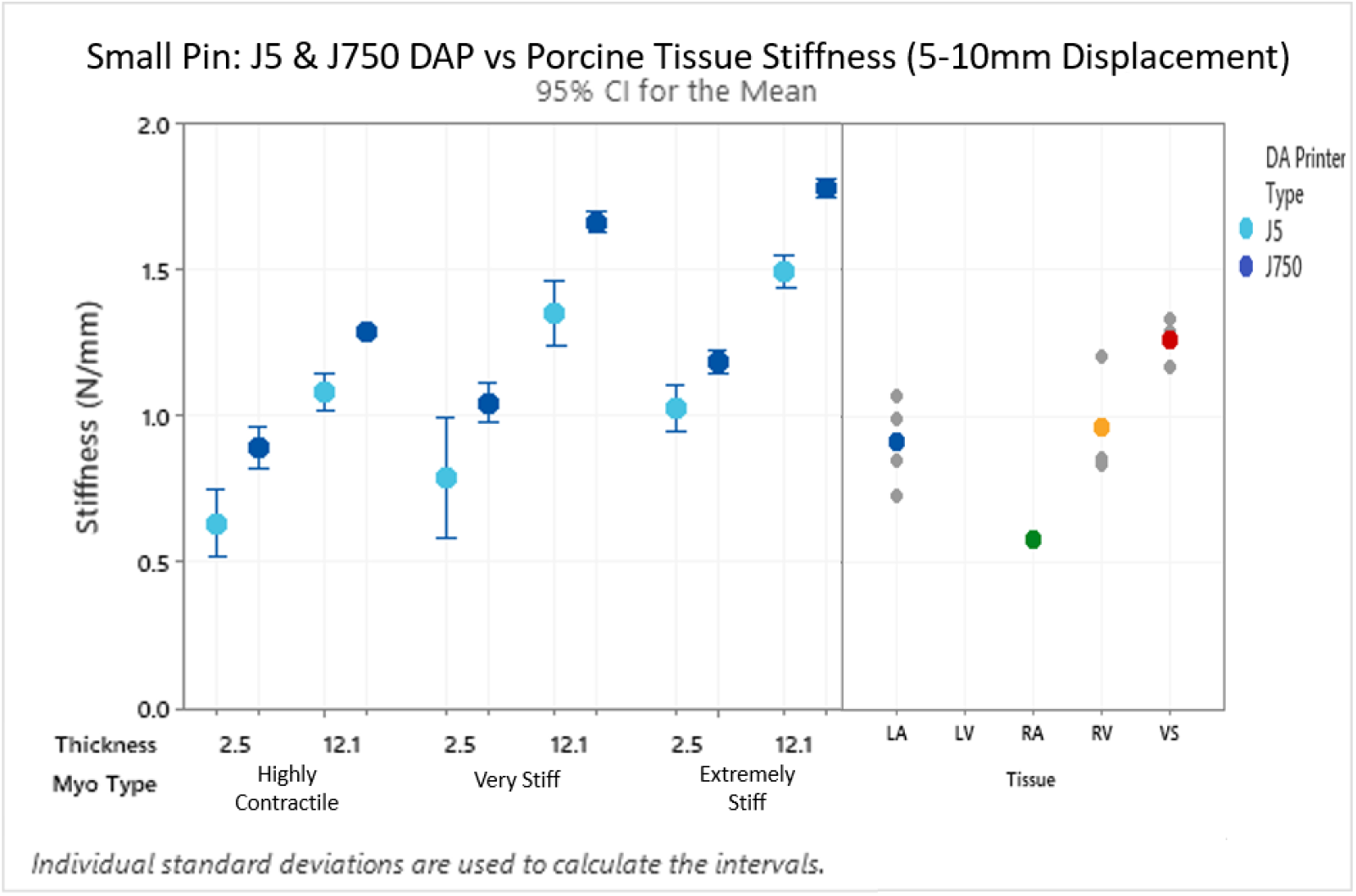
Stiffness comparison of the J5 DA Printer and J750 DA Printer myocardium samples compared to porcine tissue using a 9Fr pin displaced 10mm.

Compliance testing using a large pin displaced 5mm saw comparable stiffness values between thickness and printer types. Thicker more compliant J5 DA samples saw more variability between stiffness values compared to J750 DA samples; however, these ranges still fell within porcine tissue stiffness (Fig 10). Both printer types were observed to have higher stiffness values when compared to stiffness values that would correspond to those thicknesses within the heart.

**Figure 10.**
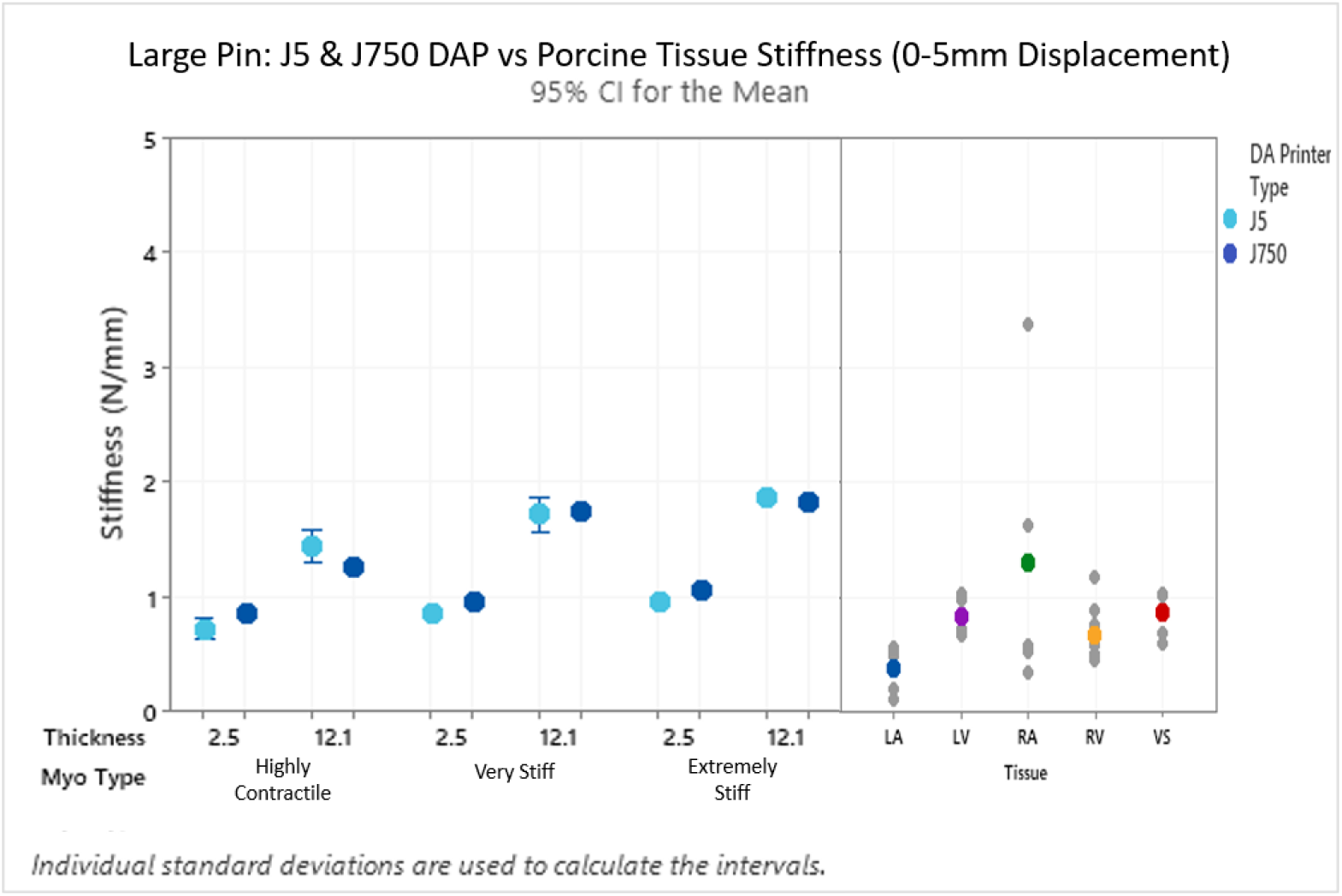
Stiffness comparison of the J5 DA Printer and J750 DA Printer myocardium samples compared to porcine tissue using a 22Fr pin displaced within the first 5mm.

Porcine samples saw high stiffness variability when displaced at larger distances. This phenomenon to a lesser degree was also observed in J5 DA printer samples especially those that are more compliant (Fig 11).

**Figure 11.**
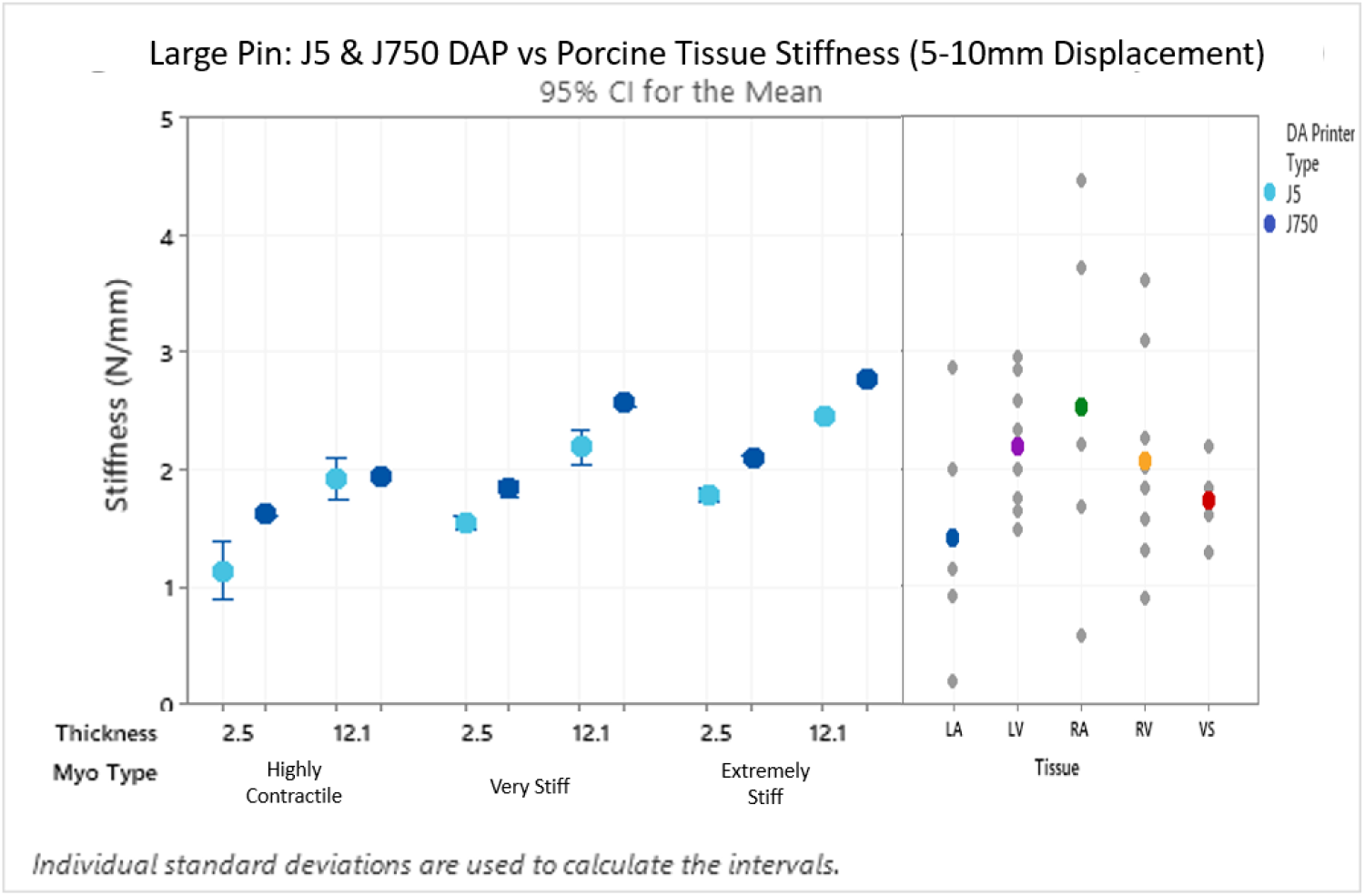
Stiffness comparison of the J5 DA Printer and J750 DA Printer myocardium samples compared to porcine tissue using a 22Fr pin displaced within the first 10mm.

### Lubricity Testing

In general, all J5 DA printer samples that were lubricated fell within the coefficient of friction ranges observed from both porcine aortic and epicardium samples. The coefficient of friction of all myocardium types between both printers produced higher values when compared to the porcine tissue samples. The samples from the J5 DA printer produced lower coefficients of friction under no lubricant testing when compared to the J750 DA samples (Fig 12a). When samples were lubricated with DI water, the coefficient of friction fell in between the tissue samples ranges. The J5 DA printer samples tended to have lower coefficient of friction when compared to the J750 DA samples and sat closer to the aortic coefficient of friction ranges (Fig 12b). Printed samples that received soap lubricant produced low coefficient of friction between both types of printers which were more closely related to aortic values (Fig 12c). Samples that received mineral oil lubricant tended to have higher coefficient of friction compared to other lubricants and were more closely related to the coefficient of friction seen from epicardium tissue values (Fig 12d). The more compliant J5 DA printer samples had lower coefficient of friction in comparison with the J750 DA printer samples and Extremely Stiff from the J5 DA printer had similar coefficient of friction when compared to the J750 DA printer Extremely Stiff samples.

**Figure 12 (a-d).**
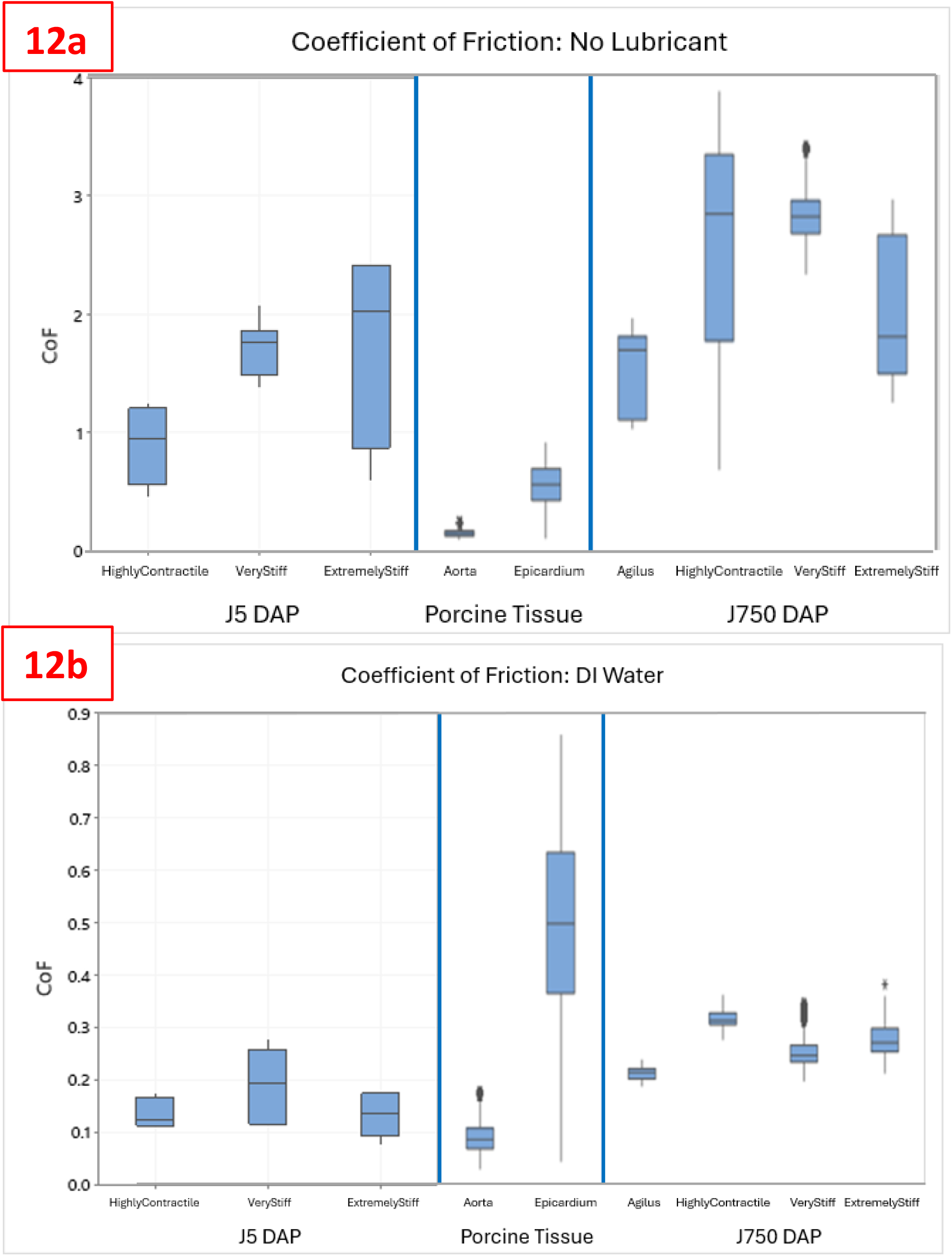
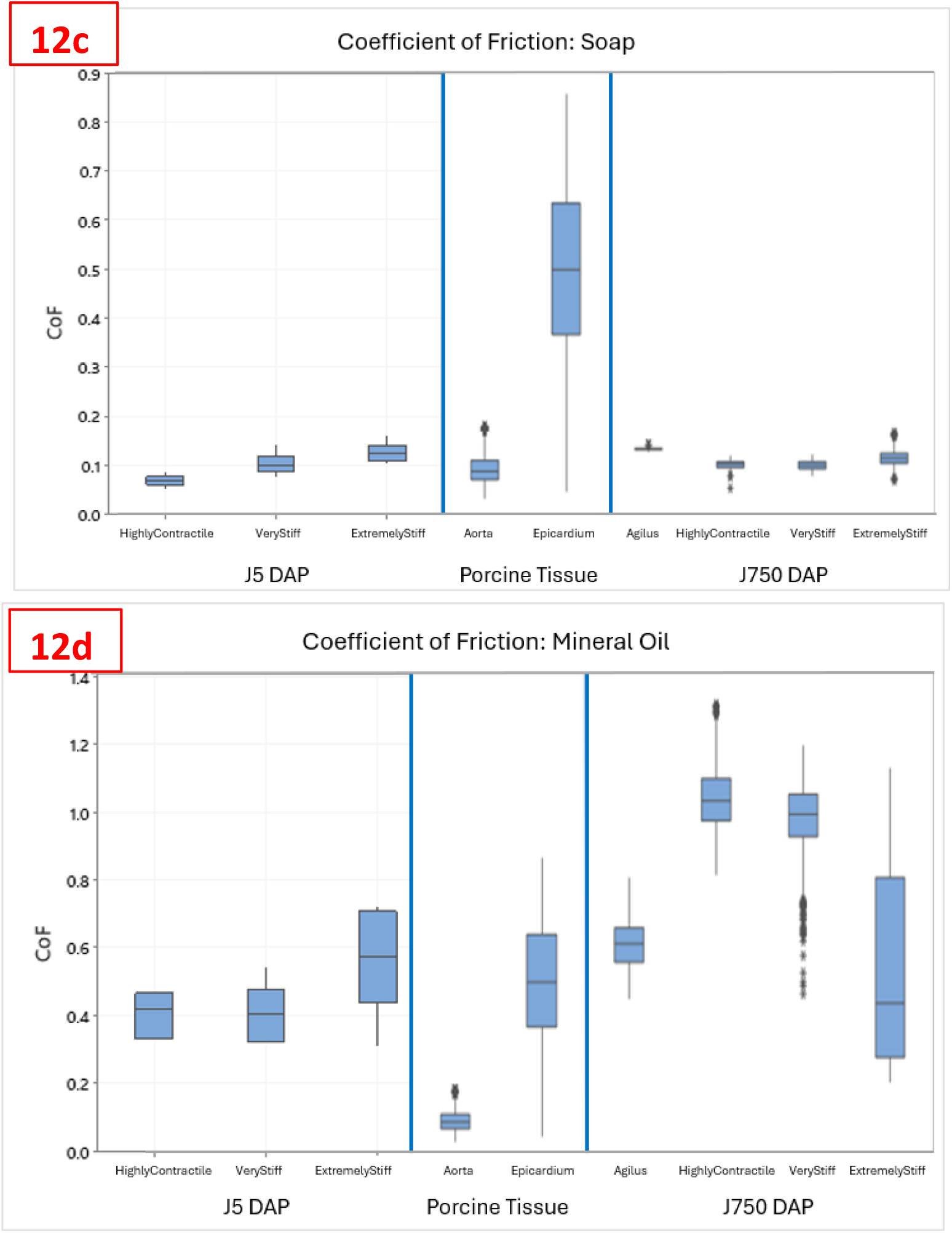
The coefficient of friction with lubricant applications: a. No Lubricant b. DI Water c. Soap and d. Mineral Oil comparison between J5 DA printed cardiac tissue sample types (left) and comparing to porcine tissue (middle) and J750 DA printed cardiac tissue samples (right)

## Discussion

Overall, this analysis compared mechanical property differences between samples from two DA printer technologies and compared them to porcine tissue. These mechanical tests included stiffness testing of compressed DA liver samples, compliance testing measuring the stiffness of DA myocardium samples, and lubricity testing to measure the coefficient of friction between DA cardiac samples. These mechanical tests were chosen to represent potential use cases between medical devices and tissue. Biological material generally contains biomechanical properties that are non-linear elasticity; however, it is observed that under small displacements there is a linear elastic portion that is captured within this testing thus making these samples ideal for certain benchtop testing [7].

### Stiffness Testing

Liver tissue is variable of thickness depending on what lobe and location of the liver is being studied, J5 DA printer liver samples saw comparable stiffnesses that was observed in porcine multi-lobe liver tissue. Overall thinner liver samples that were less compliant saw a greater difference between the two printers. It was noted previously that DA configurations are able to simulate liver tissue at low forces which correspond to small displacements and thus lower stiffness values [6]. Similar to the J750 DA printer samples, J5 DA printed liver samples also achieve this criteria thus making them acceptable to use in benchtop testing as they fall within the use conditions that would be experienced.

### Compliance Testing

Although J5 DA myocardium samples may be slightly more variable especially within thinner samples compared to the J750 DA printer, the variability is still much less when compared to porcine tissue variability. This is advantageous when conducting benchtop studies because the 3D printed tissue simulant eliminates any variability that would come from otherwise using porcine tissue.

### Lubricity Testing

Common lubricants such as DI water, soap, and mineral oil between various compliant cardiac DA samples were also found to simulate coefficient of friction values observed in aortic and epicardium porcine tissue. This then suggests that dynamic surface friction between these samples with lubricants can simulate the tactile feedback that a device may experience between these samples. Although all lubricants were found to have similar coefficients of friction to aortic and epicardium tissue, soap and DI water were found to resemble aortic surfaces while mineral oil was better suited most closely at simulating epicardium surface friction.

Overall, between all three mechanical tests conducted within this analysis, it was found that although there may be slight differences depending on the application, J5 DA samples are very similar to J750 DA printer samples and fall within the range of porcine tissue. Both printers allow for DA sample to be printed and offer different compliance of the simulated tissue which may be useful to not only simulate tissue from multiple areas of anatomy, but also to potentially simulate disease state tissues. Further analysis to include mechanical properties of disease states would be useful to further justify the usage of 3D printing in replacement of preclinical testing.

## References

[1] Pavan Kalyan BG, Kumar L. 3D Printing: Applications in Tissue Engineering, Medical Devices, and Drug Delivery. AAPS PharmSciTech. 2022 Mar 17;23(4):92. doi: 10.1208/s12249-022-02242-8. PMID: 35301602; PMCID: PMC8929713.

[2] Paul GM, Rezaienia A, Wen P, Condoor S, Parkar N, King W, Korakianitis T. Medical Applications for 3D Printing: Recent Developments. Mo Med. 2018 Jan-Feb;115(1):75–81. PMID: 30228688; PMCID: PMC6139809.

[3] Mirdamadi, E., Tashman, J. W., Shiwarski, D. J., Palchesko, R. N., & Feinberg, A. W. (2020). Fresh 3D bioprinting a full-size model of the human heart. ACS Biomaterials Science & Engineering, 6(11), 6453–6459. 10.1021/acsbiomaterials.0c01133

[4] Ma D, Gao R, Li M, Qiu J. Mechanical and medical imaging properties of 3D-printed materials as tissue equivalent materials. J Appl Clin Med Phys. 2022 Feb;23(2):e13495. doi: 10.1002/acm2.13495. Epub 2021 Dec 8. PMID: 34878729; PMCID: PMC8833282.

[5] Severseike, L., Lee, V., Brandon, T., Bakken, C., & Bhatia, V. (2019). Polyjet 3D Printing of Tissue-Mimicking Materials: How Well Can 3D Printed Synthetic Myocardium Replicate Mechanical Properties of Organic Myocardium? 10.1101/825794

[6] Lee, V., Severseike, L., Bakken, C., Bermel, E., & Bhatia, V. (2020). PolyJet 3d Printing of Tissue Mimicking Materials: An Investigation of Characteristic Properties of 3D Printed Synthetic Tissue. 10.1101/2020.12.23.424075

[7] Pinto JG, Fung YC. Mechanical properties of the heart muscle in the passive state. J Biomech. 1973 Nov;6(6):597–616. doi: 10.1016/0021-9290(73)90017-1. PMID: 4757479.

